# Genotype error biases trio-based estimates of haplotype phase accuracy

**DOI:** 10.1101/2022.04.06.487354

**Authors:** Brian L. Browning, Sharon. R. Browning

## Abstract

Haplotypes can be estimated from unphased genotype data using statistical methods. When parent-offspring data are available for inferring true phase from Mendelian inheritance rules, the accuracy of statistical phasing is usually measured by the switch error rate, which is the proportion of pairs of consecutive heterozygotes that are incorrectly phased. We present a method for estimating the genotype error rate from parent-offspring trios and a method for estimating the bias in the observed switch error rate that is caused by genotype error. We apply these methods to 485,301 genotyped UK Biobank samples that include 899 White British trios and to 38,387 sequenced TOPMed samples that include 217 African Caribbean trios and 669 European American trios. We show that genotype error inflates the observed switch error rate and that the relative bias increases with sample size. For the UK Biobank White British trios, we estimate that the observed switch error rate in the trio offspring is 2.4 times larger than the true switch error rate (1.41 × 10^−3^ vs 5.79 × 10^−4^) and that the average distance between phase errors is 64 megabases.

## Introduction

Many genetic analyses use haplotypes, which are sequences of alleles on a chromosome that are inherited from the same parent.^1-6^ Haplotypes can be inferred from unphased genotype data by means of statistical phasing methods that make use of inter-marker correlation in a sample of individuals. The accuracy of statistical phasing increases with sample size, and state of the art phasing methods can phase hundreds of thousands of genotyped individuals.^7-10^

The standard measure of phase accuracy for genome-scale data is switch error rate,^8-12^ which is the proportion of pairs of consecutive heterozygotes that are incorrectly phased. Since true phase is generally unknown, switch error rate is usually calculated using Mendelian-phased heterozygotes in trio offspring as the true phase. Mendelian phasing infers the paternally-inherited and maternally-inherited alleles in an offspring from parental genotypes and Mendelian inheritance constraints. However, genotype error can induce errors in the Mendelian phasing that lead to inflation of the observed switch error rate. In this paper, we present methods to estimate the genotype error rates and the true switch error rate.

Several methods have been developed to estimate genotype error rates. Many of these methods make use of Mendelian inheritance constraints within pedigrees. One of the first pedigree-based methods extended linkage analysis to estimate genotype error rates.^13^ This method assumes markers are in linkage equilibrium and is not designed for data with dense, correlated markers. The proportion of Mendelian inconsistent parent-offspring trios has also been used to estimate genotype error rates.^14; 15^ This approach is suitable for correlated markers, but it employs a one-parameter genotype error model that does not allow the error rate to depend on the true genotype. A recently-developed method uses inconsistent haplotype transmission in pedigrees to estimate the error rate for each combination of true and miscalled genotype.^16^ This method employs a flexible genotype error model, but it requires three generation pedigrees.

Genotype error rates can also be estimated from individuals with multiple genotyped samples.^17^ If the genotype errors in two duplicate samples are independent and if the genotype error rate is low, the genotype error rate for the individual can be estimated as one-half of the proportion of genotypes that are discordant. In reality, genotype errors in duplicate samples can be positively correlated^17^ so that one can only infer a lower bound on the genotype error rate from the genotype discordance rate. Another limitation is that the true genotype cannot be determined unless there are more than two samples for the individual (major allele homozygote, minor allele homozygote or heterozygote).

We present a likelihood-based method for estimating genotype error rates from parent-offspring trio data. Our method is suitable for dense marker data, it does not require duplicate genotypes or three generation pedigrees, and it produces an estimated genotype error rate and confidence interval for each possible true and miscalled genotype.

We apply our methods to UK Biobank SNP array data and TOPMed sequence data. We show that genotype errors in parent-offspring trios inflate the switch error rate for statistical phasing when Mendelian phased trio offspring are considered the truth. For the 485,301 UK Biobank samples, we estimate that the observed statistical phasing switch error rate in White British trio offspring is 2.4 times larger than the true rate.

## Subjects and Methods

### UK Biobank data

We filtered UK Biobank SNP array data as described previously.^10; 18^ We excluded markers with more than 5% missing genotypes, markers for which the minor allele was a singleton allele, and markers that failed one or more of the UK Biobank’s batch QC tests. We excluded samples that had been withdrawn, had an unusually high proportion of missing genotypes or heterozygous genotypes, or showed third degree or closer relationships with more than 200 individuals.^18^ After this filtering, the UK Biobank data have 711,651 autosomal markers and 487,355 samples.

We identified 1064 parent-offspring trios having 2,054 distinct parents from the pairwise kinship coefficients and discordant homozygote rates reported by the UK Biobank.^18; 19^ After excluding the 2,054 trio parents, there were 485,301 individuals. We listed the 1064 trio offspring followed by the remaining 484,237 samples in random order. We then created five subsets of the data that included 5,000, 15,000, 50,000, 150,000, and all 485,301 individuals by taking the corresponding number of samples from the head of this list. We phased each subset using Beagle 5.2.^10^

The 487,355 UK Biobank individuals include 408,938 individuals that were identified by the UK Biobank as White British based on self-report and principal component analysis. The homogenous White British subset includes 898 parent-offspring trios, which were used to estimate genotype and phase error rates in the White British individuals.

### TOPMed data

We downloaded and merged Trans-Omics for Precision Medicine (TOPMed) Program^20^ Freeze 8 high-coverage sequence data for the following studies and dbGaP^21^ accession numbers: Barbados Asthma Genetics Study (phs001143), Mount Sinai BioMe Biobank (phs001644), Cleveland Clinic Atrial Fibrillation Study (phs001189), Framingham Heart Study (phs000974), Hypertension Genetic Epidemiology Network (phs001293), Jackson Heart Study (phs000964), My Life Our Future (phs001515), Severe Asthma Research Program (phs001446), Venous Thromboembolism project (phs001402), Vanderbilt Genetic Basis of Atrial Fibrillation (phs001032), and Women’s Health Initiative (phs001237). After restricting the data to polymorphic SNVs that passed the TOPMed QC filters,^22^ there were 318,858,817 autosomal markers and 39,961 individuals.

The Barbados Asthma Genetics Study (BAGS) data include 1022 sequenced samples, and the Framingham Heart Study (FHS) data include 4166 sequenced samples. We used BAGS and FHS pedigree data to identify a set of 217 BAGS trios and a set of 669 FHS trios subject to the constraint that no individual is both a trio parent and a trio offspring. The 217 BAGS trios have 259 distinct parents, and the 669 FHS trios have 1303 distinct parents. The two sets of trios were used to estimate genotype and phase error rates in the BAGS and FHS cohorts.

After excluding all trio parents, including duplicate parent samples, there were 38,387 TOPMed samples. We listed the 886 trio offspring followed by the remaining 37,501 samples in random order. We then created four subsets of the data that included 5,000, 10,000, 20,000, and all 38,387 samples by taking the corresponding number of samples from the head of this list. We phased each subset using Beagle 5.2.^10^

### Trio genotypes

We label a marker’s major and minor alleles as A and B respectively, and we denote major allele homozygous, heterozygous, and minor allele homozygous genotypes as AA, AB, and BB respectively. When a marker has multiple alternate alleles, we aggregate all alternate alleles into a single alternate allele when applying these genotype labels. When phasing trio offspring using parental genotypes and Mendelian inheritance rules we do not aggregate alternate alleles.

At each autosomal marker, we refer to the three genotypes of a parent-offspring trio as a “trio genotype”. Two trio genotypes are considered equivalent if they each have the same set of parent genotypes and the same offspring genotypes. Note that interchanging the maternal and paternal genotypes does not change the trio genotype because this does not change the set of parental genotypes. We consider a trio genotype to be missing if any individual genotype in the trio is missing.

We express a trio genotype as a hyphen-separated list of the parent genotypes in order of increasing B-allele dose followed by the child genotype. Thus, a trio genotype with BB and AB parental genotypes and AB offspring genotype is denoted AB-BB-AB. There are 18 possible non-missing trio genotypes.

We will also express a trio genotype as a list of three B-allele doses, *x, y, z*, where *x* and *y* are the parent doses, *x* ≤ *y*, and *z* is the offspring dose. Thus, the AB-BB-AB trio genotype can also be represented as 1,2,1. We use the B-allele dose representation in superscripts and in equations.

Table 1 gives the observed autosomal trio genotype counts for the UK Biobank White British trios, the BAGS trios, and the FHS trios.

**Table 1.**
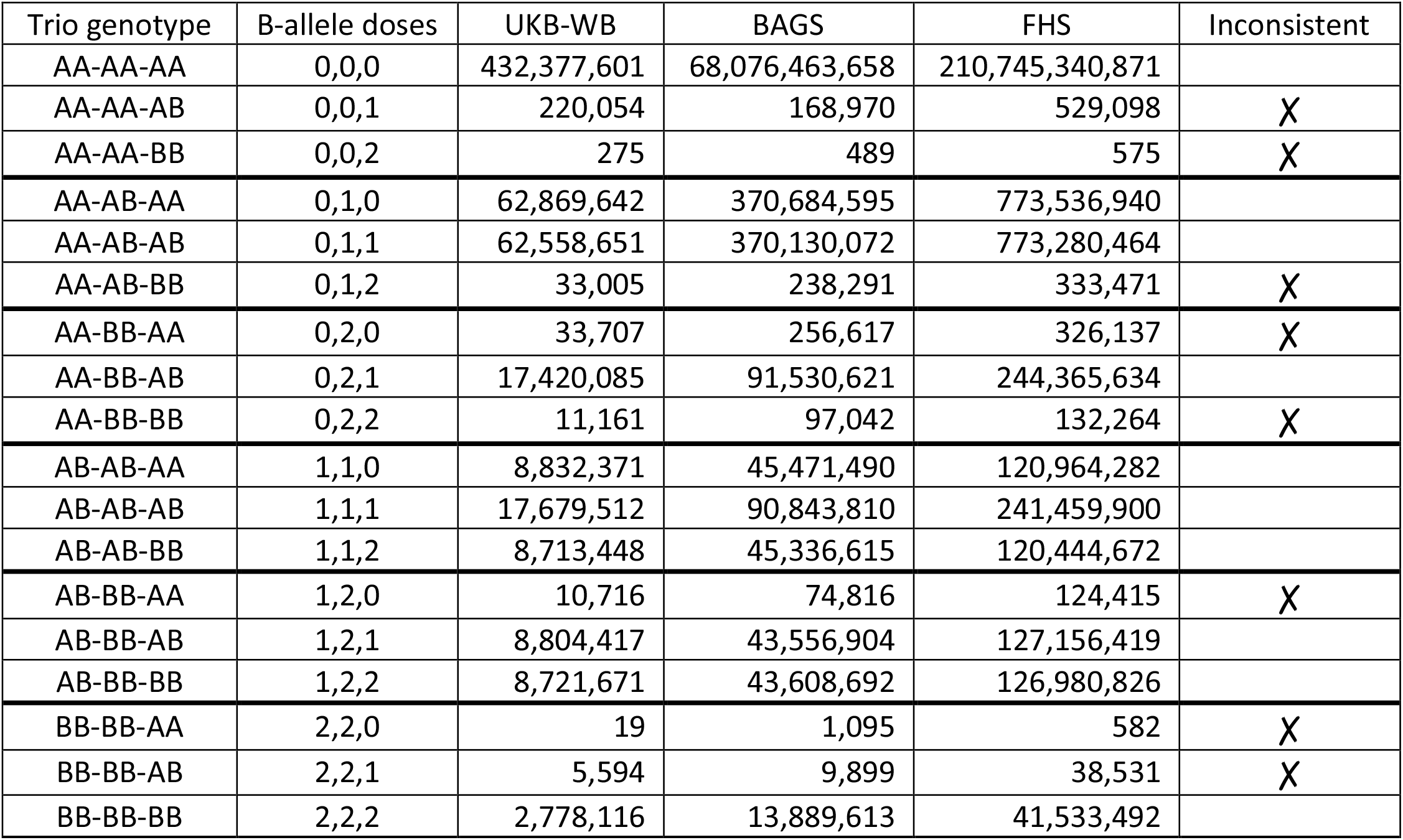
Autosomal trio genotype counts for 898 UK Biobank White British trios, 217 Barbados Asthma Genetics Study trios, and 669 Framingham Heart Study trios. A and B allele denote the major and minor allele respectively. The first two genotypes in a trio genotype are the parents’ genotypes ordered by increasing B allele dose. The last column identifies trio genotypes that are inconsistent with Mendelian inheritance rules. The UK Biobank White British trios have 7,992,553 missing trio genotypes (1.25% of trio genotypes). The TOPMed trios have no missing genotypes.

| Trio genotype | B-allele doses | UKB-WB | BAGS | FHS | Inconsistent |
| --- | --- | --- | --- | --- | --- |
| AA-AA-AA | 0,0,0 | 432,377,601 | 68,076,463,658 | 210,745,340,871 |  |
| AA-AA-AB | 0,0,1 | 220,054 | 168,970 | 529,098 | X |
| AA-AA-BB | 0,0,2 | 275 | 489 | 575 | X |
| AA-AB-AA | 0,1,0 | 62,869,642 | 370,684,595 | 773,536,940 |  |
| AA-AB-AB | 0,1,1 | 62,558,651 | 370,130,072 | 773,280,464 |  |
| AA-AB-BB | 0,1,2 | 33,005 | 238,291 | 333,471 | X |
| AA-BB-AA | 0,2,0 | 33,707 | 256,617 | 326,137 | X |
| AA-BB-AB | 0,2,1 | 17,420,085 | 91,530,621 | 244,365,634 |  |
| AA-BB-BB | 0,2,2 | 11,161 | 97,042 | 132,264 | X |
| AB-AB-AA | 1,1,0 | 8,832,371 | 45,471,490 | 120,964,282 |  |
| AB-AB-AB | 1,1,1 | 17,679,512 | 90,843,810 | 241,459,900 |  |
| AB-AB-BB | 1,1,2 | 8,713,448 | 45,336,615 | 120,444,672 |  |
| AB-BB-AA | 1,2,0 | 10,716 | 74,816 | 124,415 | X |
| AB-BB-AB | 1,2,1 | 8,804,417 | 43,556,904 | 127,156,419 |  |
| AB-BB-BB | 1,2,2 | 8,721,671 | 43,608,692 | 126,980,826 |  |
| BB-BB-AA | 2,2,0 | 19 | 1,095 | 582 | X |
| BB-BB-AB | 2,2,1 | 5,594 | 9,899 | 38,531 | X |
| BB-BB-BB | 2,2,2 | 2,778,116 | 13,889,613 | 41,533,492 |  |

### Mendelian phasing of trio offspring

If a parent has a homozygous genotype, there is only one possible allele that can be transmitted to an offspring in the absence of de novo mutation.^23^ Consequently, we can identify the paternally inherited allele and the maternally inherited allele in an offspring heterozygote if the trio genotype is non-missing and at least one parent is homozygous. We refer to heterozygous genotypes in trio offspring that have been phased using parental genotypes and Mendelian inheritance constraints as Mendelian-phased genotypes.

In this study, an offspring heterozygote is not assigned a Mendelian phase if either parent genotype is missing, if both parents are heterozygous, or if the trio genotype is inconsistent with Mendelian inheritance. The eight trio genotypes that are inconsistent with Mendelian inheritance are identified in Table 1.

### Switch error rate

The accuracy of statistical phasing is assessed using switch error rate, which is the proportion of pairs of consecutive heterozygotes that are incorrectly phased. We distinguish between the observed switch error rate and the true switch error rate. The observed switch error rate is calculated assuming the Mendelian-phasing of trio offspring is the true phase. Offspring heterozygous genotypes that do not have a Mendelian phase are ignored when calculating the observed switch error rate.

We exclude trio parents prior to statistical phasing. Consequently, the statistical phasing switch error rate in the trio offspring is an estimate of the switch error rate in individuals in the cohort who do not have a parent in the data set.

Genotype error can produce spurious, Mendelian-phased offspring heterozygotes and Mendelian phase errors. A spurious, Mendelian-phased offspring heterozygote is created whenever the true trio genotype is AA-AB-AA or AB-BB-BB and the offspring genotype is miscalled as AB. A Mendelian phase error can occur when the offspring is heterozygous and a heterozygous parental genotype is miscalled as a homozygote that does not carry the transmitted allele (Figure 1).

**Figure 1.**
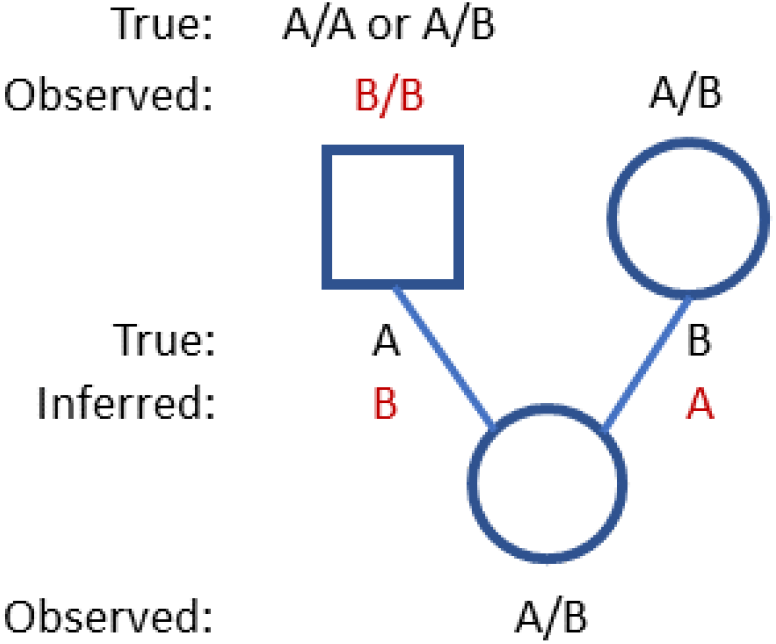
Example of a Mendelian phase error. Observed genotypes for a parent-offspring trio with alleles labeled as A and B. The father (square) transmits an A allele to the offspring and the mother (circle) transmits a B allele. If the father’s genotype is miscalled as B/B, Mendelian phasing will incorrectly infer that the offspring inherits a B allele from the father and an A allele from the mother.

De Novo mutation can also produce Mendelian phase errors. If the true trio genotype is AA-AB-AB or AB-BB-AB and a germline de novo mutation causes the AA parent to transmit a B allele or the BB parent to transmit an A allele, then the Mendelian phase of the offspring heterozygote will be incorrect. We do not specifically model Mendelian phase errors caused by de novo mutation because nearly all Mendelian phase errors are due to genotype error. A heterozygote in an offspring can be Mendelian-phased only if the site is polymorphic in the population because a parent must carry each allele. Thus, de novo mutations in the offspring can have a Mendelian phasing only if the de novo mutation occurs at a polymorphic site. The de novo mutation rate is approximately 1.3 × 10^−8^ per base pair per meiosis,^23^ and this rate is generally at least two orders of magnitude smaller than the genotype error rate at polymorphic sites.^18; 24^

### Probability of an observed trio genotype

Let 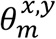 denote the probability that the true genotype at marker *m* with minor allele dose *x* ∈ {0,1,2} is called as the genotype with minor allele dose *y* ∈ {0,1,2}. The probabilities of a correct genotype call, 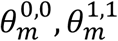, and 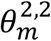, can be expressed in terms of six genotype error probabilities:

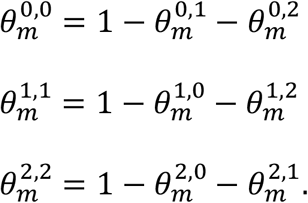

We model genotype errors as independent events that have low probability. Under this model, it would be rare for an observed trio genotype to have more than one genotype error, and this is consistent with the low frequency of AA-AA-BB and BB-BB-AA trio genotypes in Table 1.

We distinguish between true trio genotypes and observed trio genotypes. Let 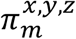 denote the population frequency of the true trio genotype *x, y, z*, at marker *m* where *x, y, z* is the B-allele dose representation of a trio genotype. Under the assumption of independent genotype errors, we can express the probability of an observed trio genotype *G*_*m*_ at marker *m* as a sum over the possible true trio genotypes. Each term of the sum is the product of the population frequency of the true trio genotype and the probability that the true genotype is called as the observed trio genotype:

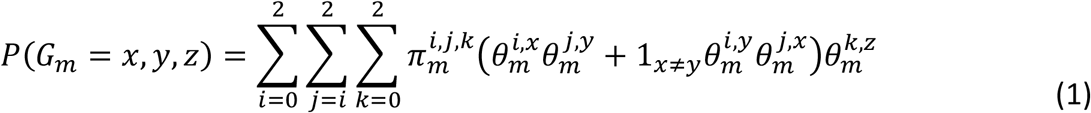

where the indicator 1_*x*≠*y*_ is 1 if *x* ≠ *y* and is 0 otherwise.

The true trio genotype frequencies 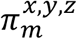 and genotype error rates 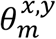 depend on the minor allele frequency (MAF). To account for this dependence, we partition the MAF spectrum (0 < MAF ≤ 0.5) into 100 subintervals of length 0.005: (0, 0.005], (0.005, 0.01], … (0.495, 0.5], and we estimate true trio genotype frequencies, 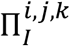, and genotype error rates, 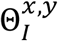, for each MAF interval *I* and for each cohort (UK Biobank White British, BAGS, and FHS).

For each cohort, we assign each marker to a MAF interval based on the marker’s observed MAF in individuals from the cohort who are not trio offspring.

### Estimating trio genotype frequencies

There are six possible sets of parental genotypes (see Table 1). For each MAF interval, we count the number of Mendelian-consistent trio genotypes in the MAF interval for each set of parental genotypes, and we estimate the probabilities of the parental genotypes to be proportional to these counts. We estimate the trio genotype frequencies 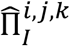 in MAF interval *I* for the *i, j, k* trio genotype from the probability of the set of parental genotypes under the assumption of transmission equilibrium.

### Estimating genotype error rates

For each MAF interval *I*, we estimate the genotype error rates 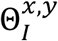 using maximum likelihood. To obtain the log likelihood, we multiply together the probabilities of the observed trio genotypes from equation (<u>1</u>) for all the non-missing trio genotypes in MAF interval *I* and take the logarithm of the product. We then replace the per-marker trio genotype frequencies 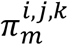 and genotype error rates 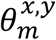 with the estimated trio genotype frequencies 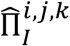and unknown genotype error rates 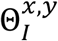 for the MAF interval. If 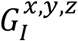 is the number of observed *x, y, z* trio genotypes at markers with MAF in interval *I*, the log likelihood function ℓ(***θ***^***I***^) for the genotype error parameters is:

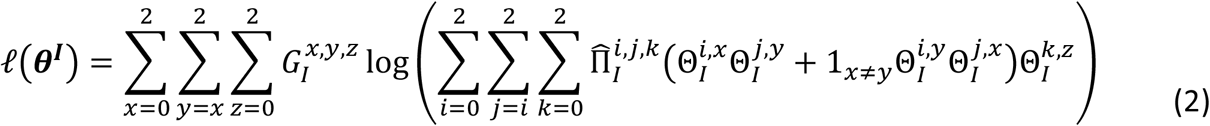

We estimate the genotype error parameters 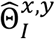by maximizing equation (<u>2</u>) using the Nelder-Mead algorithm that is implemented in the R constrOptim() function.^25^

### Estimating the number of spurious, Mendelian-phased heterozygotes

A genotype error will create a spurious, Mendelian-phased heterozygote if the true trio genotype is AA-AB-AA or AB-BB-BB and the offspring is miscalled as a heterozygote. In each MAF interval, the estimated number of spurious, Mendelian-phased heterozygotes arising from a true particular true trio genotype (AA-AB-AA or AB-BB-BB) is the product of the number of non-missing trio genotypes, the estimated trio genotype frequency and the estimated probability that the child is miscalled as a heterozygote and the parents are called correctly. We estimate the total number of spurious, Mendelian-phased heterozygotes as the sum of the estimated number in each MAF interval. See Supplemental Methods S1 for details.

### Estimating the number of Mendelian phase errors

A Mendelian phase error occurs whenever an offspring heterozygote is non-spurious and the paternally-inherited allele in the offspring is incorrectly inferred to be maternally-inherited and vice versa. There are three ways that one genotype error in a parent can create a Mendelian phase error: 1) the true trio genotype is AA-AB-AB and the AA parental genotype is miscalled as BB, 2) the true trio genotype is AB-BB-AB and the BB parental genotype is miscalled as AA, and 3) the true trio genotype is AB-AB-AB and a parental AB genotype is miscalled as a homozygous genotype that does not carry the allele transmitted to the offspring.

The estimated number of Mendelian phase errors arising from a particular true trio genotype (AA-AB-AB, AB-BB-AB, or AB-AB-AB) in a MAF interval is the product of the number of nonmissing trio genotypes, the estimated trio genotype frequency, and the estimated probability that a genotype error occurs that creates a Mendelian phase error. We estimate the number of Mendelian phase errors as the sum of the Mendelian phase errors in each MAF interval. See Supplemental Methods S2 for details.

### Estimating the true switch error rate

We derive an estimate of the true statistical phasing switch error rate for two consecutive Mendelian-phased heterozygotes by considering three cases: 1) at least one heterozygote is spurious, 2) neither heterozygote is spurious and neither or both heterozygotes have a Mendelian phase error, or 3) neither heterozygote is spurious and one heterozygote has a Mendelian phase error. From these three cases, we obtain an equation relating the observed switch error rate to the true switch error rate, the probability that a Mendelian-phased heterozygote is spurious, and the probability that a non-spurious, Mendelian-phased heterozygote has a Mendelian phase error. We use this equation to estimate the true switch error rate. See Supplemental Methods S3 for details.

### Estimating the average genotype error rate

We can estimate the average genotype error rates for markers from the estimated genotype frequencies and error rates. Since genotype error rates generally depend on allele frequency, the average genotype error rate is most interpretable for SNP arrays for which there are a fixed set of predominantly higher-frequency variants. If a study includes duplicate samples and if the duplicate genotypes are independent, the duplicate genotype discordance rate will be approximately twice the average genotype error rate.

The probability 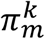 that a genotype at marker *m* has minor allele dose *k* can be expressed as the sum of frequencies of trio genotypes for which the offspring has minor allele dose *k*:

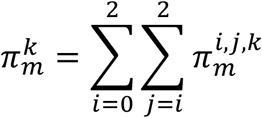

If 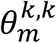 is the probability of correctly calling the genotype at marker *m*, then the probability *δ*_*m*_ of a miscalled genotype is:

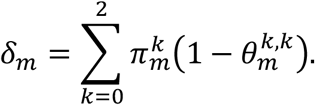

If marker *m* is in MAF interval *I*, we substitute estimated trio genotype frequencies 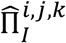 for 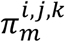 and estimated correct genotype call probabilities 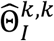 for 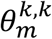. The estimated average genotype error rate is the average of the estimated genotype miscall rates, 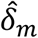.

The UK Biobank reported the genotype discordance rate for 588 pairs of experimental duplicates.^18^ In Results, we estimate the genotype error rate in the UK Biobank White British trios and compare twice this value with the reported discordance rate.

### Confidence intervals

We obtain confidence intervals for each quantity that is estimated from the observed trio genotype counts by repeating the estimation for 1000 bootstrap samples of the trios. If a data set has *t* trios, each bootstrap sample consists of *t* trios that are randomly sampled with replacement. For each set of sampled trios, we estimate the trio genotype frequencies in each MAF interval, the genotype error rates in each MAF interval, the number of spurious, Mendelian-phased heterozygotes, *S*, the number of Mendelian phase errors *M*, the true Mendelian switch error rate, *ε*_*m*_, and the true statistical phasing switch error rate, *ε*_*s*_. We obtain 95% confidence intervals for each estimated value from the 0.025 and 0.975 quantiles in the 1000 bootstrap samples.

### Single and paired switch errors

The statistical phasing of two consecutive Mendelian-phased heterozygotes will have an observed switch error if the statistical and Mendelian phasing differ. When phasing has a low error rate, it is unusual to observe more than two consecutive switch errors. Consequently, we divide switch errors into two categories: single switch errors and paired switch errors. A single switch error is a switch error that is not immediately preceded or followed by another switch error. A paired switch error is a switch error that is immediately preceded or followed by another switch error (Figure 2). We refer to two consecutive paired switch errors as a double switch error.

**Figure 2.**
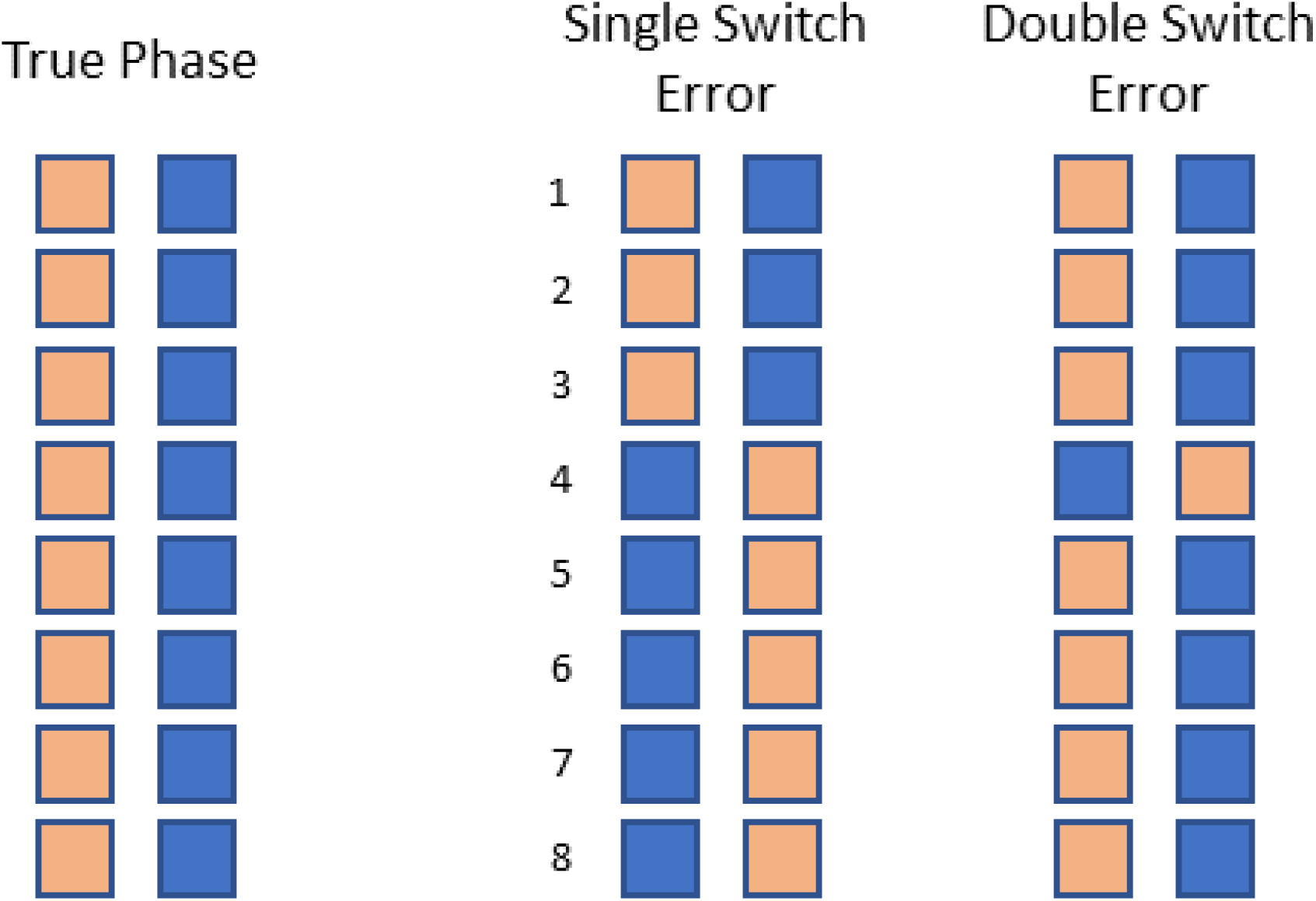
Single and double switch errors. Each column of squares represents a true or estimated haplotype at eight heterozygous genotypes. Tan and blue squares represent alleles inherited from the mother and father respectively. The left pair of columns is the true haplotype phase. A switch error occurs if a heterozygote is incorrectly phased with respect to the preceding heterozygote. The middle pair of columns has a single switch error since the 4th heterozygote is incorrectly phased with respect to the preceding heterozygote. The right pair of columns has two consecutive paired switch errors since both the 4^th^ and 5^th^ heterozygotes are incorrectly phased with respect to their preceding heterozygotes. Two consecutive paired switch errors are called a double switch error.

A single switch error indicates that the haplotypes in the individual have been incorrectly inferred in the region surrounding the switch error. In contrast, a double switch error indicates that the haplotypes have been correctly constructed in the region except at a single heterozygote whose phase with respect to surrounding heterozygotes is incorrect (Figure 2).

One disadvantage of the switch error rate as a measure of phase accuracy is that two consecutive paired switch errors are counted as two switch errors even though only one heterozygote is incorrectly phased with respect to the surrounding heterozygotes (Figure 2). To correct this deficiency, we define a phase error to be one single switch error or two consecutive paired switch errors. We can then calculate the average megabase (Mb) distance between observed phase errors as an alternate measure of phase accuracy.

We can estimate the average distance between true phase errors by accounting for Mendelian phase errors and spurious heterozygotes. If statistical phasing has a low error rate, each Mendelian phase error will introduce approximately one phase error on average, and each spurious, Mendelian-phased heterozygote will introduce approximately 0.5 phase errors on average because the statistical phasing of the spurious heterozygote is effectively random. Thus, we can estimate the true number of phase errors in statistical phasing by subtracting the estimated number of Mendelian phase errors and one-half the estimated number of spurious heterozygotes from the observed number of phase errors.

## Results

### Genotype error rates

The UK Biobank reported an average genotype discordance rate of 0.13% from 588 pairs of experimental duplicates. Since the reported genotype discordance rate was calculated without marker filtering, we applied our genotype error rate estimation method to the UK Biobank White British trio autosomes without marker filtering and estimated an average genotype error rate of 0.057% with 95% confidence interval 0.054% – 0.061%. If we assume duplicate genotypes are independent, doubling this estimated genotype error rate produces an estimated duplicate genotype discordance rate of 0.115% with 95% confidence interval 0.107% - 0.123%. This estimate is slightly less than the observed UK Biobank duplicate discordance rate of 0.13%.

We do not find any evidence of positively correlated genotype errors in the UK Biobank data. The estimated discordance rate is less than the observed discordance rate, which is the opposite of what would be expected if errors in duplicate genotype were positive correlated. The small difference in observed and estimated discordance rates may be due to the differences in the samples and markers that were used. The observed discordance was calculated from a subset of the entire UK Biobank and included X and Y chromosome genotypes and mitochondrial genotypes, whereas the estimated discordance rate was estimated from White British trios and autosomal genotypes.

After quality control filtering, we estimate an average genotype error rate of 0.054% with 95% confidence interval 0.050% – 0.058% from the UK Biobank White British trios.

Estimated genotype error rates in each MAF bin for each set of trios (UK Biobank White British, BAGS, and FHS) are shown Supplemental Figures S1, S2, and S3. These figures show that genotype error rates depend on the true genotype, with true heterozygote genotypes having the highest error rate in the sequence data.

### Observed and estimated switch error rates

Mendelian phasing statistics for each set of trios, including the average number of spurious, Mendelian-phased heterozygotes per sample and the average number of Mendelian phase errors per sample, are shown in Table 2.

**Table 2.** Average per-sample Mendelian phasing statistics for 898 UK Biobank White British trio offspring, 217 BAGS trio offspring, and 669 FHS trio offspring. Four statistics are reported for the autosomal genome in each cohort: 1) the average number of heterozygotes per sample, 2) the average number of Mendelian-phased heterozygotes per sample, 3) the estimated number of spurious, Mendelian-phased heterozygotes per sample with 95% confidence interval, and 4) the estimated number of Mendelian phase errors per sample with 95% confidence interval. Confidence intervals for each data set are obtained from 1000 bootstrap samples of the trios.

|  | Heterozygotes | Phased | Spurious Phased | Incorrectly Phased |
| --- | --- | --- | --- | --- |
| UKB-WB | 120,100 | 98,868 | 44.7 (42.2 – 48.5) | 18.7 (17.6 – 20.1) |
| BAGS | 2,747,652 | 2,328,209 | 39.8 (34.7 – 55.9) | 421.7 (403.5 – 444.2) |
| FHS | 2,072,991 | 1,711,217 | 92.4 (72.3 – 124.1) | 245.2 (225.8 – 266.0) |

Figure 3 shows the observed and estimated true switch error rates for statistical phasing as a function of sample size. The proportion of observed switch errors that are due to spurious heterozygotes and Mendelian phase errors increases with sample size. For the 485,301 UK Biobank individuals, the observed switch error rate in the White British trio offspring is 2.4 times larger than the estimated switch error (1.41 × 10^−3^ vs. 5.79 × 10^−4^).

**Figure 3.**
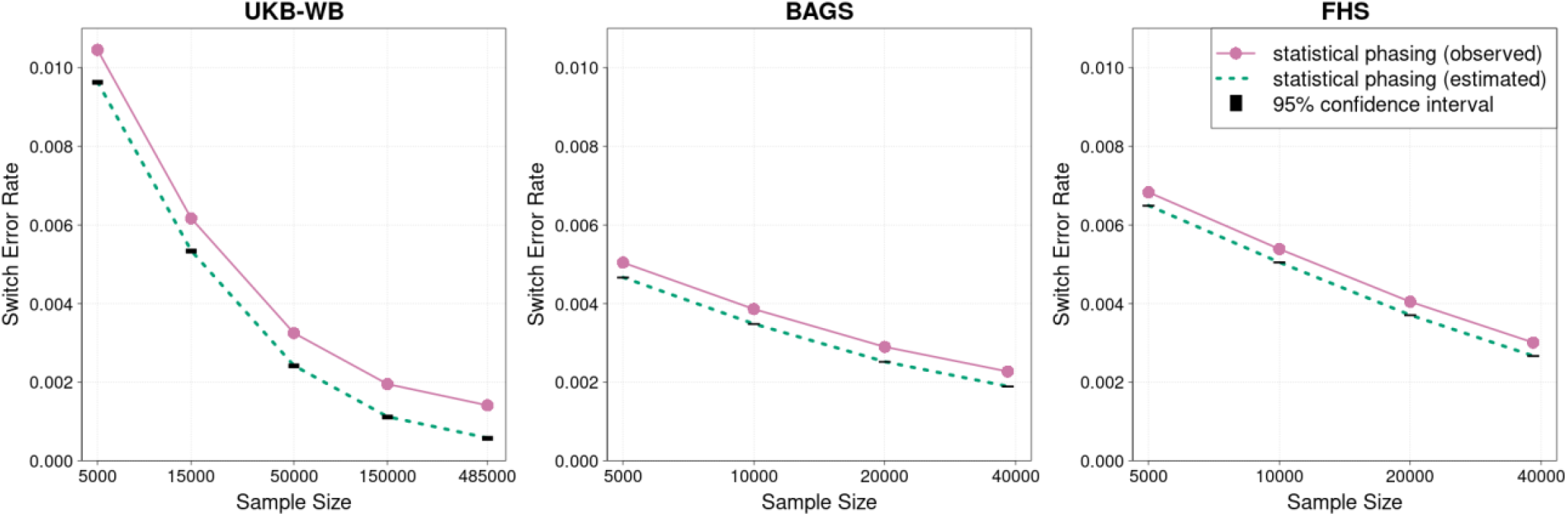
Statistical phasing switch error rates. Observed and estimated switch error rates for statistical phasing. Estimated switch error rates account for genotype errors in parent-offspring trios. The left panel shows switch error rates in 898 White British trio offspring when phasing subsets of 5,000, 15,000, 50,000, 150,000 and 485,301 individuals from the UK Biobank. The middle and right panels show switch error rates in 217 Barbados Asthma Genetic Study trio offspring (middle panel) and 669 Framingham Heart Study trio offspring (right panel) when phasing subsets of 5,000, 10,000, 20,000, and 38,387 individuals from the TOPMed study. Trio parents were excluded from the statistical phasing.

The trend of increasing relative error in the observed switch error rate as sample size increases is also observed in the TOPMed sequence data, although the maximum relative error is not as large due to the smaller number of TOPMed samples. When phasing the 38,387 TOPMed samples, the observed switch error rate in BAGS trio offspring is 20% larger than the estimated switch error rate (2.27 × 10^−3^ vs 1.89 × 10^−3^), and the observed switch error rate in the FHS trio offspring is 13% larger than the estimated switch error rate (3.01 × 10^−3^ vs 2.67 × 10^−3^).

### Average distance between phase errors

When phasing the 485,301 UK Biobank individuals, 79% of observed switch errors in the White British trio offspring are paired switch errors. When phasing the 38,387 TOPMed samples, 94% and 92% of observed switch errors are paired switch errors in the BAGS and FHS trio offspring respectively.

Table 3 shows average distance between observed switch errors, the average distance between observed phase errors, and the estimated average distance between true phase errors for each set of trio offspring. Average distance is calculated as the ratio of the Mb distance spanned by the autosomal markers to the number of switch or phase errors.

**Table 3.** Average megabases between autosomal phasing errors for 898 White British trio offspring in 485,301 statistically-phased UK Biobank individuals, and for 217 Barbados Asthma Genetic Study trio offspring and 669 Framingham Heart Study trio offspring in 38,387 statistically-phased TOPMed samples. Trio parents are excluded from the statistical phasing. A phase error is a single switch error or a double switch error.

|  | Observed Mb / switch error | Observed Mb / phase error | Estimated Mb / phase error |
| --- | --- | --- | --- |
| UKB-WB | 20.05 | 33.10 | 64.54 |
| BAGS | 0.53 | 1.00 | 1.18 |
| FHS | 0.54 | 1.01 | 1.13 |

## Discussion

We have presented a method for estimating the genotype error rate from parent-offspring trios and a method for estimating the statistical phasing switch error rate that accounts for spurious heterozygotes and Mendelian phase errors. The method for estimating the genotype error rate does not require duplicate genotypes, and it produces an estimated error rate stratified by MAF interval for each possible true and miscalled genotype (major allele homozygote, minor allele homozygote, or heterozygote). We applied these methods to 898 White British trios included in 485,301 genotyped UK Biobank individuals and to 217 African Caribbean trios and 669 European American trios included in 38,387 samples sequenced by the TOPMed project.

We found that the relative inflation in the observed switch error rate due to genotype error increases as sample size increases. For the 485,301 UK Biobank samples, the observed switch error rate in White British trio offspring is 2.4 times larger than the estimated switch error rate. Inflation in the observed switch error rate is also seen in the TOPMed sequence data, but the relative inflation in the TOPMed data is less than in the UK Biobank data. This difference is primarily due to the smaller TOPMed sample size.

Most observed switch errors in these data are paired switch errors. Two consecutive paired switch errors indicate that the two haplotypes in a region have been constructed correctly except for one incorrectly phased heterozygote (Figure 2). Since two consecutive paired switch errors represent a single incorrectly phased heterozygote, we recommend counting two consecutive paired switch errors as a single phasing error when reporting the average distance between phase errors. With this definition of phase error, we estimate that there is an average of 43 phase errors per UK Biobank White British trio offspring in the autosomal genome (Table 3), which is less than two phase errors per chromosome. For comparison, we estimate that there are an average of 45 spurious heterozygotes per chromosome in these same individuals (Table 2).

Our methods require a set of parent-offspring trios from a homogenous population. For the UK Biobank, we applied our methods to the subset of White British individuals, since this is the largest genetically homogenous subset in the UK Biobank. The observed switch error rate in White British trio offspring is lower than the observed switch error rate in the other 166 trio offspring (0.00141 vs 0.00333) because the White British subset has a larger sample size and less genetic heterogeneity.

Our analysis of sequence data is limited to SNVs. In principle, the methods presented here can be extended to other types of variants, but genotype error rates for copy number variants and more complex structural variants may be higher. Estimation of these other error rates is an area for future research.

Subsets of heterozygotes on a chromosome can also be phased using information from sequence reads.^26^ The phase of any two heterozygotes in the same phased set is inferred with respect to each other, but the phase of two heterozygotes from different phased sets is not, and some heterozygotes may be left unphased. However, sequence reads have the potential to phase the heterozygotes that can only be phased with 50% accuracy using statistical methods, such as singleton variants. A recent evaluation showed that incorporating phase information from sequence reads in statistical phasing can achieve higher accuracy than is possible with either statistical phasing or read-based phasing by itself.^9^ However, incorporating phase information from sequence read data for a large number of samples is computationally challenging.

Parent-offspring trios are a valuable resource for assessing data accuracy. Trios have been the primary tool for assessing statistical phase accuracy. This study shows that trios can also be used to estimate genotype error rates, and that genotype accuracy must be considered when estimating the phase error rate in large-scale data.

## Data and code availability

UK Biobank data were downloaded from the European Genome-Phenome Archive (https://ega-archive.org/datasets/EGAD00010001497). TOPMed freeze 8 data for the following study accession numbers were downloaded from dbGaP (https://www.ncbi.nlm.nih.gov/gap/): phs001143, phs001644, phs001189, phs000974, phs001293, phs000964, phs001515, phs001446, phs001402, phs001032, phs001237.

## Supplemental Data

Supplemental Data includes supplemental methods, three figures, and supplemental acknowledgements

## Acknowledgements

Research reported in this publication was supported by the National Human Genome Research Institute of the National Institutes of Health under award number HG008359. The content is solely the responsibility of the authors and does not necessarily represent the official views of the National Institutes of Health.

This research has been conducted using the UK Biobank Resource under Application Number 19934. Molecular data for the Trans-Omics in Precision Medicine (TOPMed) program was supported by the National Heart, Lung and Blood Institute (NHLBI). Core support including centralized genomic read mapping and genotype calling, along with variant quality metrics and filtering were provided by the TOPMed Informatics Research Center (3R01HL-117626-02S1; contract HHSN268201800002I). Core support including phenotype harmonization, data management, sample-identity QC, and general program coordination were provided by the TOPMed Data Coordinating Center (R01HL-120393; U01HL-120393; contract HHSN268201800001I). We gratefully acknowledge the studies and participants who provided biological samples and data for TOPMed. Funding for the Barbados Asthma Genetics Study was provided by National Institutes of Health (NIH) R01HL104608, R01HL087699, and HL104608 S1. The Framingham Heart Study was supported by contracts NO1-HC-25195, HHSN268201500001I and 75N92019D00031 from the NHLBI and grant supplement R01 HL092577-06S1; genome sequencing was funded by HHSN268201600034I and U54HG003067. See Supplemental Data for acknowledgments of additional studies in the TOPMed data.

## Supplemental Data

Methods S1: Estimating the number of spurious, Mendelian-phased heterozygotes

Methods S2: Estimating the number of Mendelian phase errors

Methods S3: Estimating the true statistical phasing switch error rate

Figure S1: UK Biobank White British genotype error rates stratified by minor allele frequency

Figure S2: TOPMed BAGS genotype error rates stratified by minor allele frequency

Figure S3: TOPMed FHS genotype error rates stratified by minor allele frequency Supplemental Acknowledgements

## Supplemental Methods

### Methods S1: Estimating the number of spurious, Mendelian-phased heterozygotes

A genotype error will create a spurious, Mendelian-phased heterozygote if the true trio genotype is AA-AB-AA or AB-BB-BB and the offspring is miscalled as a heterozygote.

Let *n*_*I*_ be the number of non-missing trio genotypes in MAF interval *I*, and let 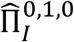 and 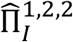 respectively be the estimated trio genotype frequencies for the AA-AB-AA and AB-BB-BB trio genotypes in MAF interval *I* (see “Estimating trio genotype frequencies” in Methods). We estimate the number of true AA-AB-AA trio genotypes as 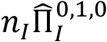, and we estimate the number of true AB-BB-BB trio genotypes as 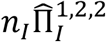. Let 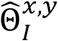 be the estimated probability that a genotype with minor-allele dose *x* ∈ {0,1,2} is called as a genotype with minor-allele dose *y* ∈ {0,1,2} (see “Estimating genotype error rates” in Methods).

In each MAF interval, the estimated number of spurious, Mendelian-phased heterozygotes arising from a true AA-AB-AA or AB-BB-BB trio genotype is the product of the estimated true trio genotype count and the estimated probability that the child is miscalled as a heterozygote and the parents are called correctly.

The estimated number of spurious, Mendelian-phased heterozygotes *Ŝ* is the sum of the estimated counts in each MAF interval:

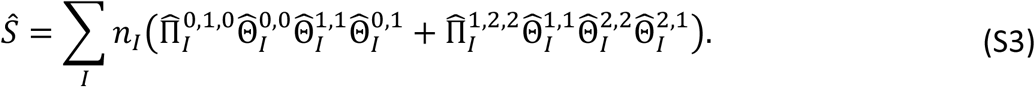

### Methods S2: Estimating the number of Mendelian phase errors

A Mendelian phase error occurs whenever an offspring heterozygote is non-spurious and Mendelian phasing incorrectly infers that the paternally-inherited allele in the offspring is maternally-inherited and vice versa. There are three ways that one genotype error can create a Mendelian phase error: 1) the true trio genotype is AA-AB-AB and the AA parental genotype is miscalled as BB, 2) the true trio genotype is AB-BB-AB and the BB parental genotype is miscalled as AA, and 3) the true trio genotype is AB-AB-AB and one parental AB genotype is miscalled as a homozygous genotype that does not carry the allele transmitted to the offspring.

Let *n*_*I*_ be the number of non-missing trio genotypes in MAF interval *I*, and let 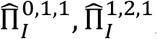, and 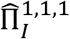 respectively be the estimated trio genotype frequencies for the AA-AB-AB, AB-BB-AB, or AB-AB-AB trio genotypes in MAF interval *I* (see “Estimating trio genotype frequencies” in Methods). We estimate the number of true AA-AB-AB trio genotypes as 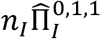, we estimate the number of true AB-BB-AB trio genotypes as 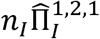, and we estimate the number of true AB-AB-AB trio genotypes as 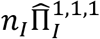. Let 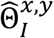 be the estimated probability that a genotype with minor-allele dose *x* ∈ {0,1,2} is called as a genotype with minor-allele dose *y* ∈ {0,1,2} (see “Estimating genotype error rates” in Methods).

In each MAF interval, the estimated number of Mendelian phase errors is the sum of the Mendelian phase errors in the three error cases. We estimated number of true AA-AB-AB trio genotypes in MAF interval *I* for which the AA parental genotype is miscalled as BB as 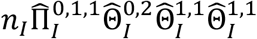. We estimate the number of true AB-BB-AB trio genotypes in MAF interval *I* for which the BB parental genotype is miscalled as AA as 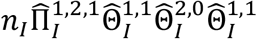.

In the third error case, the true trio genotype is AB-AB-AB and one parental AB genotype is miscalled as a homozygote that does not carry the transmitted allele. In MAF interval *I*, we estimate the probability that one parental AB genotype in the trio is miscalled as a homozygote as 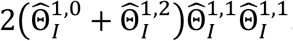. If a parental AB genotype is miscalled as a homozygote, the probability that the erroneous homozygote does not carry the transmitted allele is 0.5 because the A and B allele have equal probability of being transmitted to the offspring. Thus, we estimate the number of true AB-AB-AB in MAF interval *I* for which one AB parental genotype is miscalled as a homozygote that does not carry the transmitted allele as

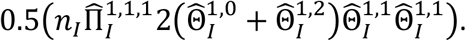

The estimated number of Mendelian phase errors 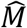 is the sum of the estimated Mendelian phase errors in each MAF interval:

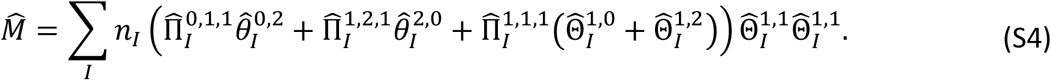

### Methods S3: Estimating the true statistical phasing switch error rate

We derive an expression for the true statistical phasing switch error rate for two consecutive Mendelian-phased heterozygotes by considering three cases: 1) at least one heterozygote is spurious, 2) neither heterozygote is spurious and neither or both heterozygotes have a Mendelian phase error, or 3) neither heterozygote is spurious and one heterozygote has a Mendelian phase error.

In the following cases, *p*_*S*_ is the probability that a Mendelian-phased heterozygote is spurious, *p*_*M*_ is the probability that a non-spurious, Mendelian-phased heterozygote has a phase error, and *ε*_*s*_ is the true statistical phasing switch error rate for two consecutive non-spurious, Mendelian-phased heterozygotes.

Case 1. The probability that at least one of two consecutive Mendelian-phased heterozygotes is spurious is 1 − (1 − *p*_*S*_)^2^. In this case, the statistical phasing of the two heterozygotes will differ from the Mendelian phasing with probability 0.5 because the statistical phasing is effectively random.

Case 2. The probability that neither heterozygote is spurious neither or both heterozygotes have a Mendelian phase error is 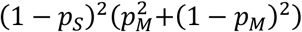. In this case the Mendelian phasing of the two heterozygotes does not have a switch error and the statistical phasing of the two heterozygotes will differ from the Mendelian phasing with probability *ε*_*s*_.

Case 3. The probability that neither heterozygote is spurious and that one heterozygote has a Mendelian phase error is (1 − *p*_*S*_)^2^(2*p*_*M*_(1 − *p*_*M*_)). In this case the Mendelian phasing of the two heterozygotes has a switch error, and the statistical phasing of the two heterozygotes will differ from the Mendelian phasing with probability 1 − *ε*_*s*_.

Combining the probabilities for the preceding three cases, we obtain the following equation for the expected observed switch error rate *ε*_0_ for two consecutive Mendelian-phased heterozygotes:

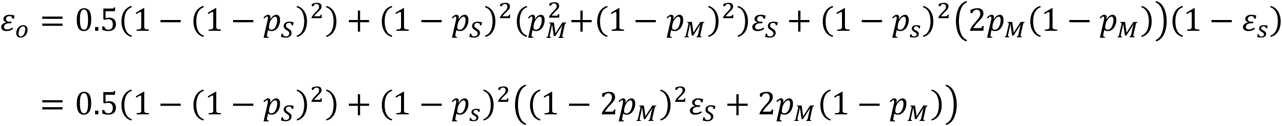

Solving the preceding equation for the true statistical phasing switch error rate *ε*_*s*_ gives

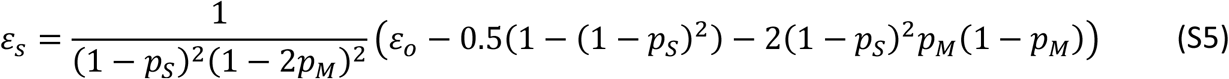

We estimate *p*_*S*_ as 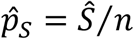 where *Ŝ* is the estimated number of spurious, Mendelian-phased heterozygotes from equation (S1) and *n* is the number of Mendelian-phased heterozygotes. We estimate *p*_*M*_ as 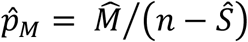 where 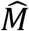 is the estimated number of Mendelian phase errors from equation (S2). Substituting the observed switch error rate for *ε*_0_ and the estimates 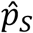 and 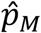 into equation (<u>S5</u>) gives an estimate 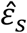 of the true statistical phasing switch error rate.

## Supplemental Figures

**Figure S1.**
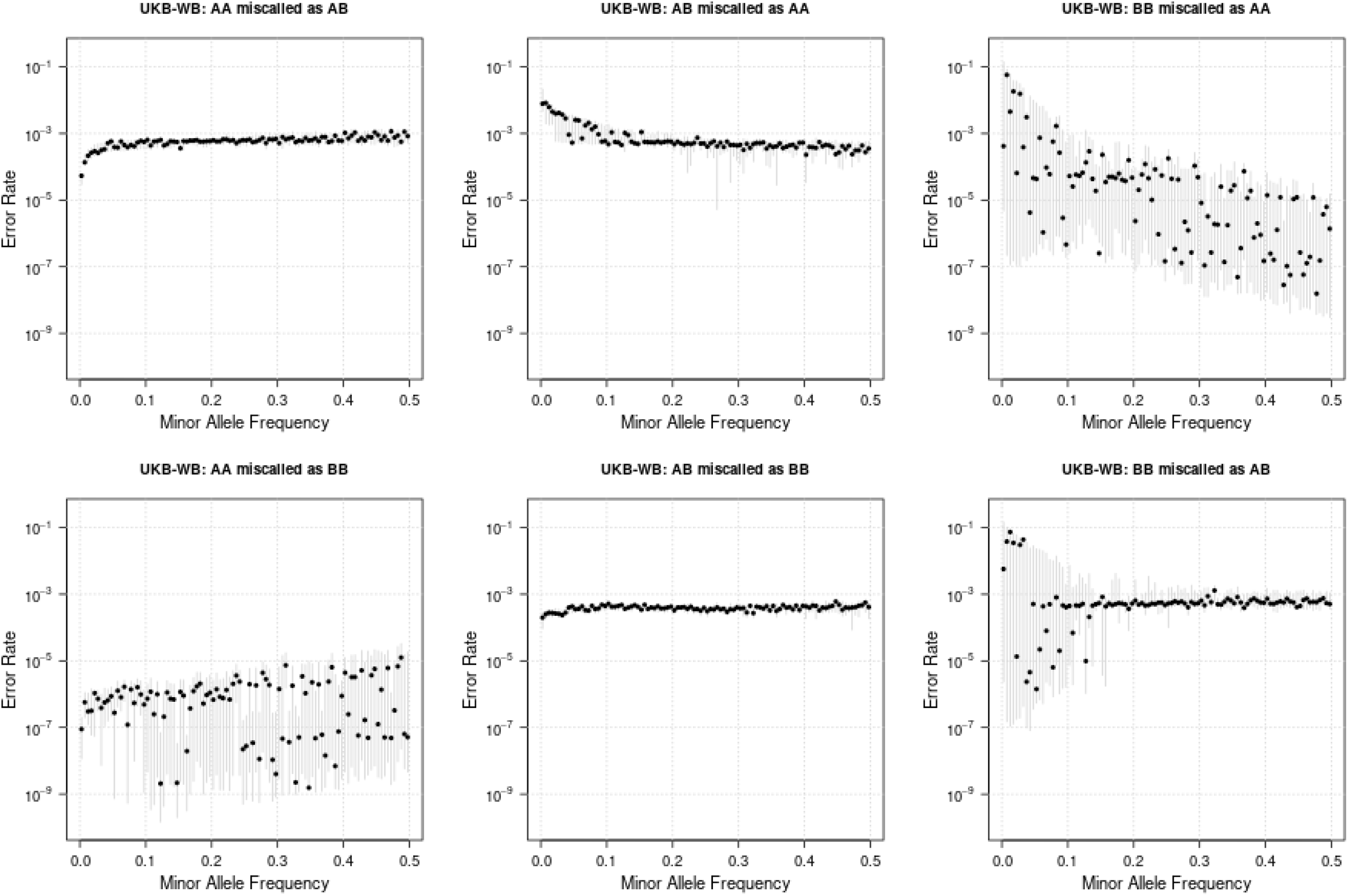
Estimated genotype error rates stratified by minor allele frequency (MAF) in 898 UK Biobank White British (UKB-WB) parent-offspring trios. Homozygous major allele, heterozygous, and homozygous minor allele genotypes are denoted AA, AB, and BB respectively. The MAF spectrum (0 < MAF ≤ 0.5) is partitioned into 100 subintervals of length 0.005, and markers are assigned to MAF intervals corresponding to their observed MAF in 408,040 White British individuals who are not trio offspring. The six panels show estimated genotype error rates for each minor allele frequency interval. Shaded vertical lines are 95% bootstrap confidence intervals calculated by sampling trios with replacement.

**Figure S2.**
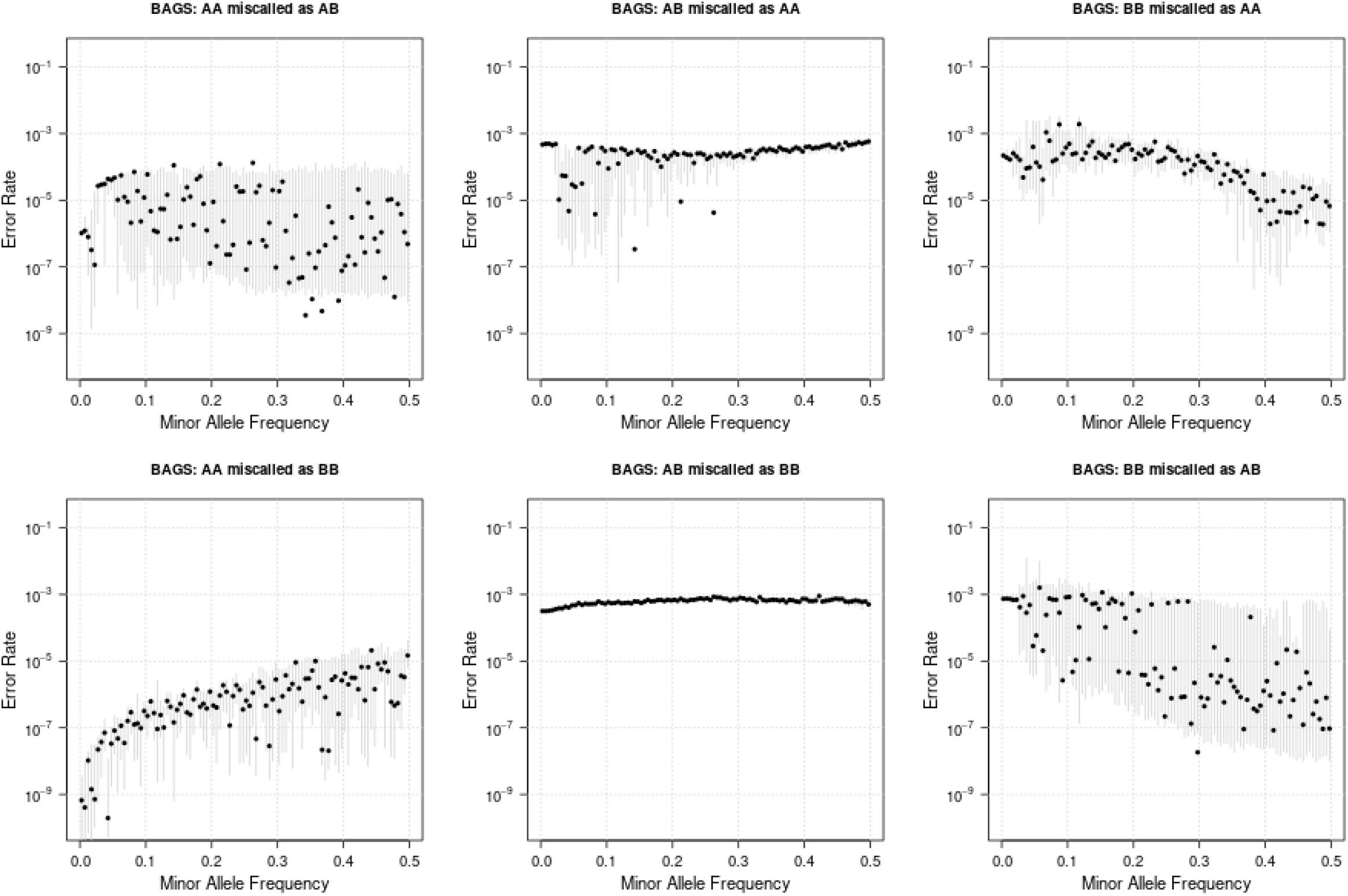
Estimated genotype error rates stratified by minor allele frequency (MAF) in 217 parent-offspring trios from the Barbados Asthma Genetic Study (BAGS). Homozygous major allele, heterozygous, and homozygous minor allele genotypes are denoted AA, AB, and BB respectively. The MAF spectrum (0 < MAF ≤ 0.5) is partitioned into 100 subintervals of length 0.005, and markers are assigned to MAF intervals corresponding to their observed MAF in 786 BAGS individuals who are not trio offspring. The six panels show estimated genotype error rates for each minor allele frequency interval. Shaded vertical lines are 95% bootstrap confidence intervals calculated by sampling trios with replacement.

**Figure S3.**
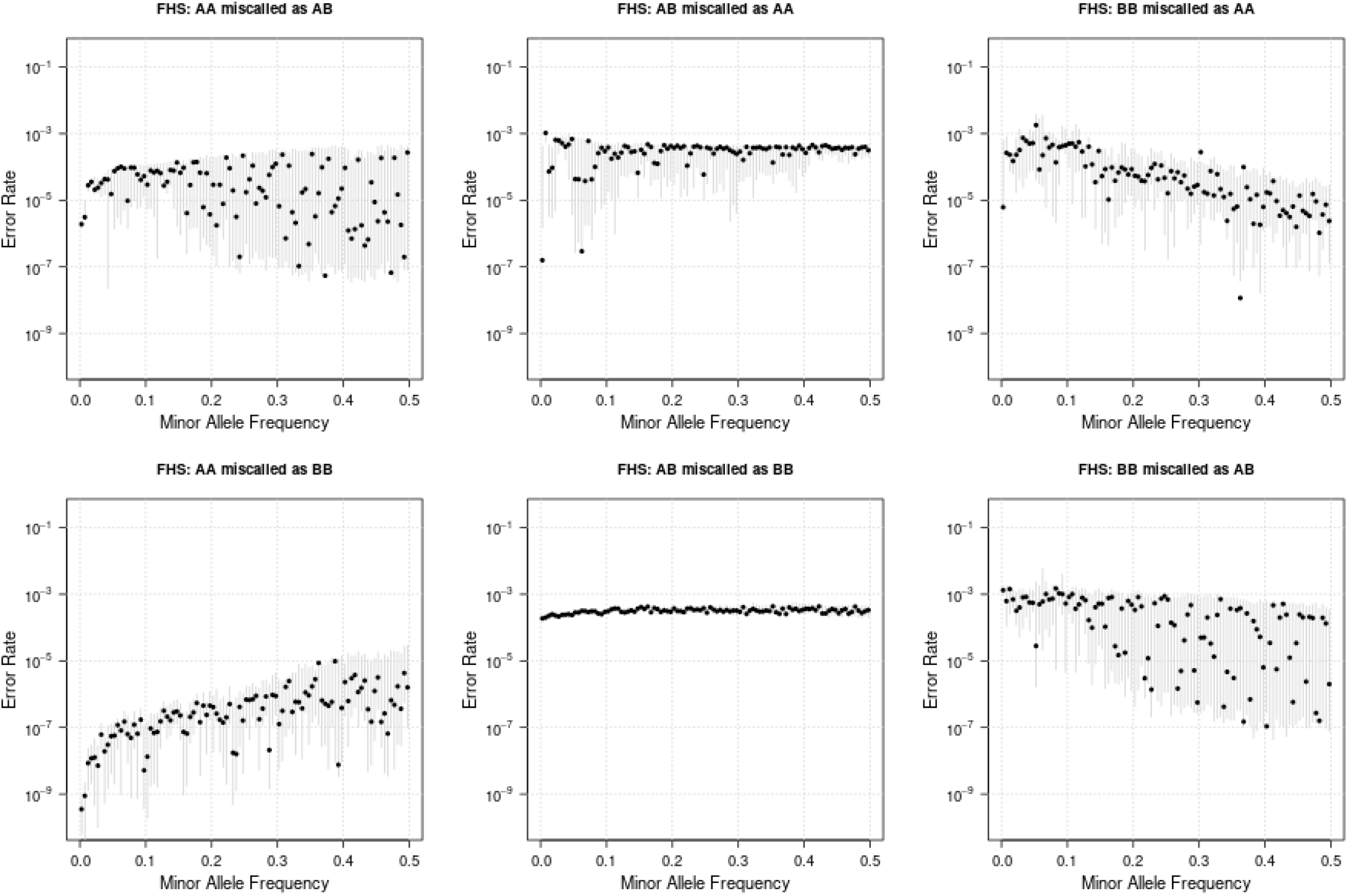
Estimated genotype error rates stratified by minor allele frequency (MAF) in 669 parent-offspring trios from the Framingham Heart Study (FHS). Homozygous major allele, heterozygous, and homozygous minor allele genotypes are denoted AA, AB, and BB respectively. The MAF spectrum (0 < MAF ≤ 0.5) is partitioned into 100 subintervals of length 0.005, and markers are assigned to MAF intervals corresponding to their observed MAF in 3,485 FHS individuals who are not trio offspring. The six panels show estimated genotype error rates for each minor allele frequency interval. Shaded vertical lines are 95% bootstrap confidence intervals calculated by sampling trios with replacement.

## Supplemental Acknowledgments

We gratefully acknowledge the studies and participants who provided biological samples and data for TOPMed. Funding for the Barbados Asthma Genetics Study was provided by National Institutes of Health (NIH) R01HL104608, R01HL087699, and HL104608 S1. The Mount Sinai BioMe Biobank has been supported by The Andrea and Charles Bronfman Philanthropies and in part by funds from the NHLBI and the National Human Genome Research Institute (NHGRI) (U01HG00638001; U01HG007417; X01HL134588); genome sequencing was funded by contract HHSN268201600037I. The Cleveland Clinic Atrial Fibrillation study was supported by NIH grants R01 HL 090620 and R01 HL 111314, the NIH National Center for Research Resources for Case Western Reserve University and Cleveland Clinic Clinical and Translational Science Award UL1-RR024989, the Cleveland Clinic Department of Cardiovascular Medicine philanthropy research funds, and the Tomsich Atrial Fibrillation Research Fund; genome sequencing was supported by R01HL092577. The Framingham Heart Study was supported by contracts NO1-HC-25195, HHSN268201500001I and 75N92019D00031 from the NHLBI and grant supplement R01 HL092577-06S1; genome sequencing was funded by HHSN268201600034I and U54HG003067. The Hypertension Genetic Epidemiology Network Study is part of the NHLBI Family Blood Pressure Program; collection of the data represented here was supported by grants U01 HL054472, U01 HL054473, U01 HL054495, and U01 HL054509; genome sequencing was funded by R01HL055673. The Jackson Heart Study is supported and conducted in collaboration with Jackson State University (HHSN268201300049C and HHSN268201300050C), Tougaloo College (HHSN268201300048C), and the University of Mississippi Medical Center (HHSN268201300046C and HHSN268201300047C) contracts from NHLBI and the National Institute for Minority Health and Health Disparities (NIMHD); genome sequencing was funded by HHSN268201100037C. The My Life, Our Future samples and data are made possible through the partnership of Bloodworks Northwest, the American Thrombosis and Hemostasis Network, the National Hemophilia Foundation, and Bioverativ; genome sequencing was funded by HHSN268201600033I and HHSN268201500016C. The Severe Asthma Research Program was conducted with the support of the NHLBI grants R01 HL069116, R01 HL069130, R01 HL069149, R01 HL069155, R01 HL069167, R01 HL069170, R01 HL069174, R01 HL069349, U10 HL109086, U10 HL109146, U10 HL109152, U10 HL109164, U10 HL109168, U10 HL109172, U10 HL109250, and U10 HL109257; genome sequencing was funded by HHSN268201500016C. The Venous Thromboembolism project was funded in part by grants from the NIH, NHLBI (HL66216 and HL83141) and the NHGRI (HG04735). The Vanderbilt Genetic Basis of Atrial Fibrillation study was supported by grants from the American Heart Association (EIA 0940116N), and grants from the National Institutes of Health (HL092217, U19 HL65962, and UL1 RR024975), and by CTSA award (UL1TR000445) from the National Center for Advancing Translational Sciences; genome sequencing was funded by R01HL092577. The Women’s Health Initiative program is funded by NHLBI through contracts 75N92021D00001, 75N92021D00002, 75N92021D00003, 75N92021D00004, 75N92021D00005; genome sequencing was funded by HHSN268201500014C.

## References

1. Howie, B., Fuchsberger, C., Stephens, M., Marchini, J., and Abecasis, G.R. (2012). Fast and accurate genotype imputation in genome-wide association studies through pre-phasing. Nat Genet 44, 955–959.

2. Browning, S.R., and Browning, B.L. (2020). Probabilistic Estimation of Identity by Descent Segment Endpoints and Detection of Recent Selection. Am J Hum Genet 107, 895–910.

3. Ramstetter, M.D., Dyer, T.D., Lehman, D.M., Curran, J.E., Duggirala, R., Blangero, J., Mezey, J.G., and Williams, A.L. (2017). Benchmarking relatedness inference methods with genome-wide data from thousands of relatives. Genetics 207, 75–82.

4. Zhou, Y., Browning, B.L., and Browning, S. (2019). Population-specific recombination maps from segments of identity by descent. bioRxiv, 868091.

5. Zhou, Y., Browning, S.R., and Browning, B.L. (2020). A Fast and Simple Method for Detecting Identity-by-Descent Segments in Large-Scale Data. Am J Hum Genet 106, 426–437.

6. Maples, B.K., Gravel, S., Kenny, E.E., and Bustamante, C.D. (2013). RFMix: a discriminative modeling approach for rapid and robust local-ancestry inference. Am J Hum Genet 93, 278–288.

7. Loh, P.R., Palamara, P.F., and Price, A.L. (2016). Fast and accurate long-range phasing in a UK Biobank cohort. Nat Genet 48, 811–816.

8. Loh, P.-R., Danecek, P., Palamara, P.F., Fuchsberger, C., Reshef, Y.A., Finucane, H.K., Schoenherr, S., Forer, L., McCarthy, S., and Abecasis, G.R. (2016). Reference-based phasing using the Haplotype Reference Consortium panel. Nat Genet 48, 1443.

9. Delaneau, O., Zagury, J.F., Robinson, M.R., Marchini, J.L., and Dermitzakis, E.T. (2019). Accurate, scalable and integrative haplotype estimation. Nat Commun 10, 5436.

10. Browning, B.L., Tian, X., Zhou, Y., and Browning, S.R. (2021). Fast two-stage phasing of large-scale sequence data. Am J Hum Genet 108, 1880–1890.

11. Li, Y., Willer, C.J., Ding, J., Scheet, P., and Abecasis, G.R. (2010). MaCH: using sequence and genotype data to estimate haplotypes and unobserved genotypes. Genet Epidemiol 34, 816–834.

12. Williams, A.L., Patterson, N., Glessner, J., Hakonarson, H., and Reich, D. (2012). Phasing of many thousands of genotyped samples. Am J Hum Genet 91, 238–251.

13. Sobel, E., Papp, J.C., and Lange, K. (2002). Detection and integration of genotyping errors in statistical genetics. Am J Hum Genet 70, 496–508.

14. Hao, K., Li, C., Rosenow, C., and Hung Wong, W. (2004). Estimation of genotype error rate using samples with pedigree information--an application on the GeneChip Mapping 10K array. Genomics 84, 623–630.

15. Saunders, I.W., Brohede, J., and Hannan, G.N. (2007). Estimating genotyping error rates from Mendelian errors in SNP array genotypes and their impact on inference. Genomics 90, 291–296.

16. Wang, R.J., Radivojac, P., and Hahn, M.W. (2021). Distinct error rates for reference and nonreference genotypes estimated by pedigree analysis. Genetics 217, 1–10.

17. Wall, J.D., Tang, L.F., Zerbe, B., Kvale, M.N., Kwok, P.Y., Schaefer, C., and Risch, N. (2014). Estimating genotype error rates from high-coverage next-generation sequence data. Genome Res 24, 1734–1739.

18. Bycroft, C., Freeman, C., Petkova, D., Band, G., Elliott, L.T., Sharp, K., Motyer, A., Vukcevic, D., Delaneau, O., O’Connell, J., et al. (2018). The UK Biobank resource with deep phenotyping and genomic data. Nature 562, 203–209.

19. Manichaikul, A., Mychaleckyj, J.C., Rich, S.S., Daly, K., Sale, M., and Chen, W.-M. (2010). Robust relationship inference in genome-wide association studies. Bioinformatics 26, 2867–2873.

20. Taliun, D., Harris, D.N., Kessler, M.D., Carlson, J., Szpiech, Z.A., Torres, R., Taliun, S.A.G., Corvelo, A., Gogarten, S.M., and Kang, H.M. (2019). Sequencing of 53,831 diverse genomes from the NHLBI TOPMed Program. bioRxiv, 563866.

21. Mailman, M.D., Feolo, M., Jin, Y., Kimura, M., Tryka, K., Bagoutdinov, R., Hao, L., Kiang, A., Paschall, J., Phan, L., et al. (2007). The NCBI dbGaP database of genotypes and phenotypes. Nat Genet 39, 1181–1186.

22. Danecek, P., Auton, A., Abecasis, G., Albers, C.A., Banks, E., Depristo, M.A., Handsaker, R.E., Lunter, G., Marth, G.T., Sherry, S.T., et al. (2011). The variant call format and VCFtools. Bioinformatics 27, 2156–2158.

23. Tian, X., Browning, B.L., and Browning, S.R. (2019). Estimating the Genome-wide Mutation Rate with Three-Way Identity by Descent. Am J Hum Genet 105, 883–893.

24. Taliun, D., Harris, D.N., Kessler, M.D., Carlson, J., Szpiech, Z.A., Torres, R., Taliun, S.A.G., Corvelo, A., Gogarten, S.M., Kang, H.M., et al. (2021). Sequencing of 53,831 diverse genomes from the NHLBI TOPMed Program. Nature 590, 290–299.

25. R Development Core Team. (2006). R: A Language and Environment for Statistical Computing.(Vienna, Austria: R Foundation for Statistical Computing).

26. Garg, S. (2021). Computational methods for chromosome-scale haplotype reconstruction. Genome Biol 22, 101.

